# PARP1 and MGMT interaction-based sensitivity to DNA damage in Ewing sarcoma

**DOI:** 10.1101/2020.01.26.920405

**Authors:** Dauren Alimbetov, Jodie Cropper, Rostislav Likhotvorik, Ruth Carlson, Youngho Kwon, Raushan Kurmasheva

**Affiliations:** Greehey Children’s Cancer Research Institute, University of Texas Health at San Antonio, San Antonio, TX 78229, USA; Department of Molecular Medicine, University of Texas Health at San Antonio, San Antonio, TX 78229, USA; Department of Biochemistry and Structural Biology, University of Texas Health at San Antonio, San Antonio, TX 78229, USA

**Keywords:** protein-protein interaction, DNA damage, DNA repair, DNA methyltransferase, base excision repair (BER), inhibition mechanism, cancer therapy, drug resistance

## Abstract

The Ewing family of sarcomas comprises the fourth most common highly aggressive bone tumor. Four of five Ewing sarcoma chemotherapeutics induce DNA damage, as does radiation therapy. At relapse, two additional DNA-damaging agents are routinely used to re-induce remission, indicating that Ewing sarcoma is intrinsically sensitive to DNA damage. However, current treatment regimens are relatively ineffective, specifically for relapsed or metastatic disease. Several preclinical studies, including our study in the Pediatric Preclinical Testing Program (PPTP), provide evidence for the synthetic lethal combination of PARP1 inhibitor talazoparib with a DNA-methylating agent, temozolomide, for Ewing sarcoma. Nevertheless, in both preclinical studies and clinical trials, doses of temozolomide were significantly reduced because of toxicity of the drug combination. Temozolomide-induced DNA lesions are repaired *via* poly(ADP) ribose polymerase I (PARP1)-dependent base excision repair and by O6-methylguanine-DNA methyltransferase (MGMT) in a single-step adduct removal. Here, we provide evidence that the two DNA repair pathways act in an epistatic manner in lesion removal. Further, we demonstrate that PARP1 and MGMT physically interact, and that this association is stimulated upon DNA damage. Protein co-immunoprecipitation and microscale thermophoresis analyses revealed that PARP1/MGMT complex formation is DNA and PARylation-independent. Collectively, our results show that: 1) DNA damage response pathways mediated by PARP1 and MGMT work epistatically to eliminate temozolomide-induced DNA adducts; 2) PARP1 and MGMT physically interact; and 3) PARP1/MGMT interaction is increased in response to DNA damage. We discuss how our findings may affect therapeutic advancement for Ewing sarcoma and potentially other cancer types.

---

Ewing sarcoma is a bone or soft tissue malignancy affecting adolescents and young adults, with particularly poor outcomes for patients with metastatic or relapsed disease. Cells are intrinsically sensitive to DNA-damaging agents (1-3). Current standard chemotherapy for Ewing sarcoma includes a combination of vincristine, doxorubicin, and cyclophosphamide, alternating with etoposode and ifosfamide, plus radiation (4-9). Four of these drugs induce DNA damage, as does radiation therapy. About 80% of relapse occurs within 2 years of initial diagnosis; when this occurs, two additional DNA-damaging agents (irinotecan and temozolomide (TMZ)) are routinely added to re-induce remission (10-12). European protocols generally combine vincristine, doxorubicin, and an alkylating agent in a single treatment cycle (13). Unlike regional or local disease, this therapy is relatively ineffective for metastatic tumors, with event-free survival in only 12% of cases and overall long-term survival in 9% to 33% of cases (14,15).

Ewing sarcoma is driven by a fusion oncogene, EWSR1-FLI1, resulting from a reciprocal translocation event involving EWSR1 (chromosome 22) and the ETS family transcription factor FLI1 (chromosome 11). Poly-ADP ribose polymerase 1 (PARP1) is a direct transcriptional target of EWSR1-FLI1 (16,17). The fusion oncogene induces high levels of RNAPII transcription while BRCA1 is sequestered with this active transcription complex, conferring a homologous recombination defect (18). Therefore, it has been postulated that inhibiting PARP1 activity *via* inhibitors selectively impacts Ewing sarcoma cells, either by downregulating activity of the oncogenic EWSR1-FLI1 fusion protein or due to the homologous recombination defect of these cells (19,20). Our earlier results in the PPTP (Pediatric Preclinical Testing Program) demonstrated that as a single agent, the PARP1 inhibitor talazoparib (TLZ) has only modest activity in *in vivo* pediatric xenograft tumor models without defects in homologous recombination, although it has promising activity *in vitro* (17). However, our testing of the TLZ/TMZ combination showed significant synergy in 5 of 10 Ewing sarcoma xenograft models (21). Consistent with our preclinical studies(21), high toxicity of this combination was also observed in patients (22), which required a reduced (and potentially sub-optimal) dose of TLZ. These findings emphasize the need for developing new treatment regimens for Ewing sarcoma (23).

The synthetic lethality induced by PARP1 inhibitor and DNA-damaging agents, such as TMZ, has been reported by several groups (24,25). However, the mechanism of this synergistic cytotoxicity, and the basis for their selective action specifically in Ewing sarcoma, remains unclear. Two current predominant models postulate that either inhibition of PARP1 catalytic activity or PARP1 trapping on DNA leads to cell death in the presence of TMZ-induced DNA lesions (26). Indeed, the DNA adduct(s) responsible for synergistic killing have not been defined.

PARP1 is an abundant nuclear protein that catalyzes the post-translational modification of many cellular targets *via* i.e. poly(ADP) ribosylation or PARylation, and works in concert with O_6_-methylguanine methyltransferase (MGMT) to repair damage through base excision repair or DNA mismatch repair (27-29). TMZ acts as a prodrug that is converted to the active metabolite 5-(3-methyltriazen-1-yl) imidazole-4-carboxamide, which induces predominantly methyl adducts at N3 of adenine (N3-meA), and N7 and O6 of guanine (N7-meG, O6-meG) in DNA (29-31). N3-meA and N7-meG adducts are repaired *via* PARP1 in BER, while TMZ toxicity is primarily mediated through the O6-meG, which is highly cyto- and genotoxic. The O6-meG adducts are removed through a suicidal mechanism of MGMT (32-36); in MGMT-deficient cells, an unrepaired O6-meG mispairs with thymine but not cytosine during the DNA replication that leads to activation of MMR (37-39). Therefore, low levels of MGMT warrant a good response to TMZ. Thus, modulation of MGMT and PARP1 activity is an effective approach to overcoming cellular resistance to TMZ, and enhancing response to the drug combination. In this study, we find that PARP1 and MGMT act in an epistatic manner to repair TMZ-generated DNA adducts, and demonstrate that this functional cooperation may stem from a direct PARP1/MGMT protein interaction, which has not been shown before (33) (schematic view shown in **Fig. 1**). Altogether, our results highlight the possibility of targeting the PARP1/MGMT complex in development of new and better treatment regimens for Ewing sarcoma.

**Figure 1.**
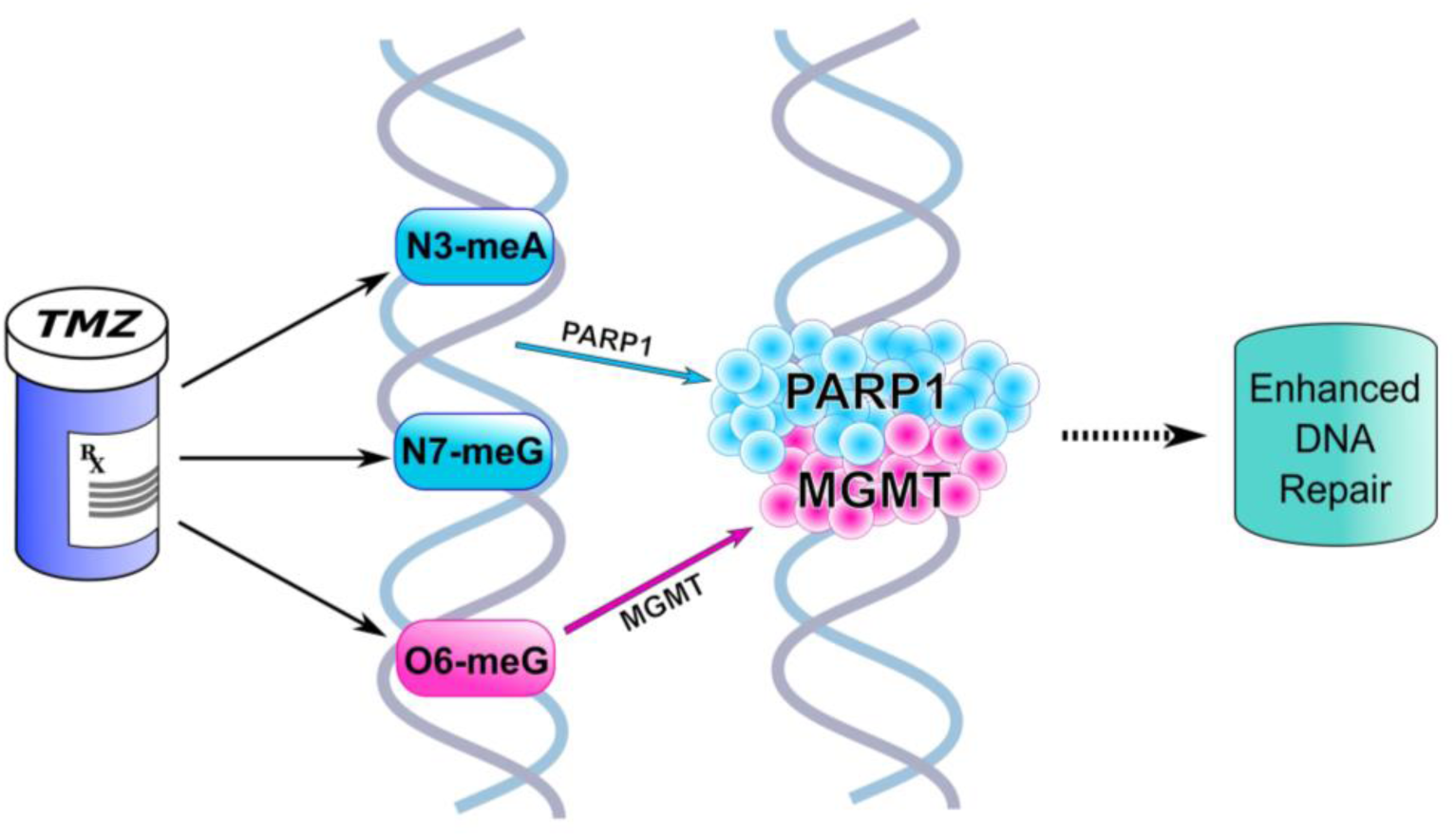
Schematic view of the proposed TMZ-induced DNA damage and repair in Ewing sarcoma. TMZ induces three major methyl-adducts on DNA (N3-meA, N7-meG, and O6-meG), which are repaired by either PARP1 in BER or by MGMT in one-step removal of methyl-group (29). PARP1 and MGMT physically interact in response to DNA damage induction by TMZ and enhance DNA repair. The image was created with *Inkscape* and *GNU Image Manipulation Program* graphic packages.

## Results

### PARP1 and MGMT function epistatically to eliminate TMZ-generated adducts

Because of the role of MGMT in removal of the O6-meG adduct induced by TMZ (29), we investigated whether MGMT contributes to TMZ resistance in Ewing sarcoma cells. Previously, we investigated the intrinsic sensistivity of Ewing sarcoma cells to TLZ and TMZ (**Table 1**). Next, two cell lines either sensitive (ES-4, ES-7) or resistant (EW-8, ES-6) to TLZ and TMZ were exposed to TMZ, TLZ, and O6-benzylguanine (MGMT inhibitor). When cells were treated with TMZ+TLZ, TMZ+O6-benzylguanine, or TMZ+TLZ+O6-benzylguanine, cell sensitivity to TMZ was essentially similar (**Fig. 2**). Thus, O6-benzylguanine fully sensitized cells, and addition of TLZ did not further enhance sensitivity. Conversely, O6-benzylguanine did not further enhance cell sensitivity to TMZ in the presence of TLZ. These effects were observed in all four cell lines. Since the most toxic lesion is O6-meG, these results suggest that sensitization to TMZ is mediated by a pathway *epistatic* with removal of O6-meG, demonstrating that this toxic adduct is repaired by a pathway using both MGMT and PARP1. This mechanism also implies that TLZ sensitizes Ewing sarcoma cells by inhibiting repair of O6-meG, not being trapped on DNA in response to N3-meA and N7-meG adducts.

**Table 1.** Intrinsic sensitivity of Ewing sarcoma cell lines to TLZ and TMZ.

| **Response** | **Cell line** | **TLZ**<br>**IC<sub>10</sub> (nM)** | **TMZ**<br>**IC<sub>50</sub> (μM)** |
| --- | --- | --- | --- |
| **Sensitive** | ES-4 | 1.6 | 112.0 |
|  | ES-7 | 0.7 | 446.3 |
| **Resistant** | ES-6 | 7.2 | 12008.0 |
|  | EW-8 | 29.7 | 1436.0 |

TLZ, talazoparib; TMZ, temozolomide

**Figure 2.**
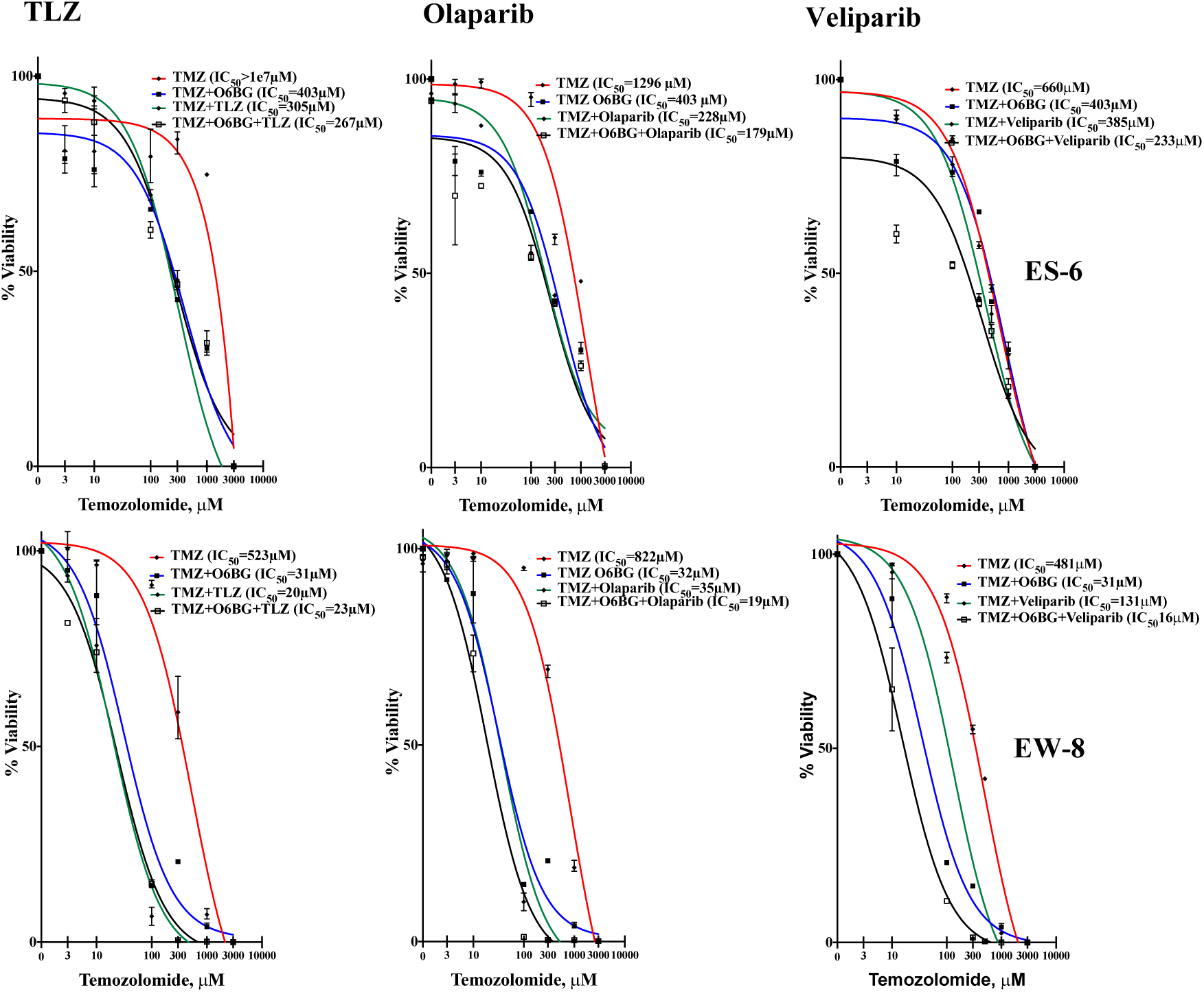
Epistatic potentiation of TMZ in ES-6 and EW-8 cells is not enhanced by TLZ, olaparib, or veliparib over that achieved by O6-benzylguanine. TMZ-treated cells (0-3mM) were exposed to a PARP inhibitor (IC_10_) and O6-benzylguanine (5μM) for 96hrs (Alamar Blue staining). Red curve, TMZ; blue, (TMZ + O6-benzylguanine); green, TMZ + (TLZ/olaparib/veliparib); black TMZ + O6-benzylguanine + (TLZ/olaparib/veliparib).

To determine whether the DNA trapping potency of a PARP1 ihibitor could contribute to sensitizing cells to TMZ, we investigated the effects of TLZ (high potency), olaparib (medium potency), and veliparib (low potency). TLZ is the most potent DNA-trapping PARP1 inhibitor now in clinical development; its DNA trapping potency is ∼100-fold greater than olaparib and ∼500-fold greater than veliparib (40). It is shown that catalytic activity is inversely correlated with DNA trapping potency in these inhibitors. All three PARP1 inhibitors sensitized cells to TMZ to a similar extent, suggesting that the ability of PARP1 ihibitors to sensitize cells to TMZ (which requires repair of O6-meG), is independent of the PARP1 DNA trapping potency of the inhibitor **(Fig.2)**. Thus, these data demonstrate that PARP1 activity is required for removal of O6-meG adducts.

### PARP1 and MGMT proteins physically interact in DNA damage repair

The epistatic mechanism of DNA repair by PARP1 and MGMT implies that these proteins cooperate to provide enhanced function for DNA damage repair. To test this conjecture, we performed PARP1/MGMT co-immunoprecipitation in EW-8 cells because the epistatic response to TLZ and TMZ showed no dependence on cell type. Both PARP1 and MGMT immunoprecipitation resulted in co-precipitating PARP1 or MGMT proteins by immunoblotting, which indicated that PARP1 and MGMT exist in a complex. Higher levels of MGMT and PARP1 proteins were detected in the complex from TMZ-treated cells (**Fig. 3A** and **B**, lane 2) compared to untreated cells (lane 1). The increased protein signal in lane 2 indicates that in response to DNA damage, MGMT and PARP1 tightly associate to repair DNA adducts, and they are maintained at lower amounts when stress levels are absent.

**Figure 3.**
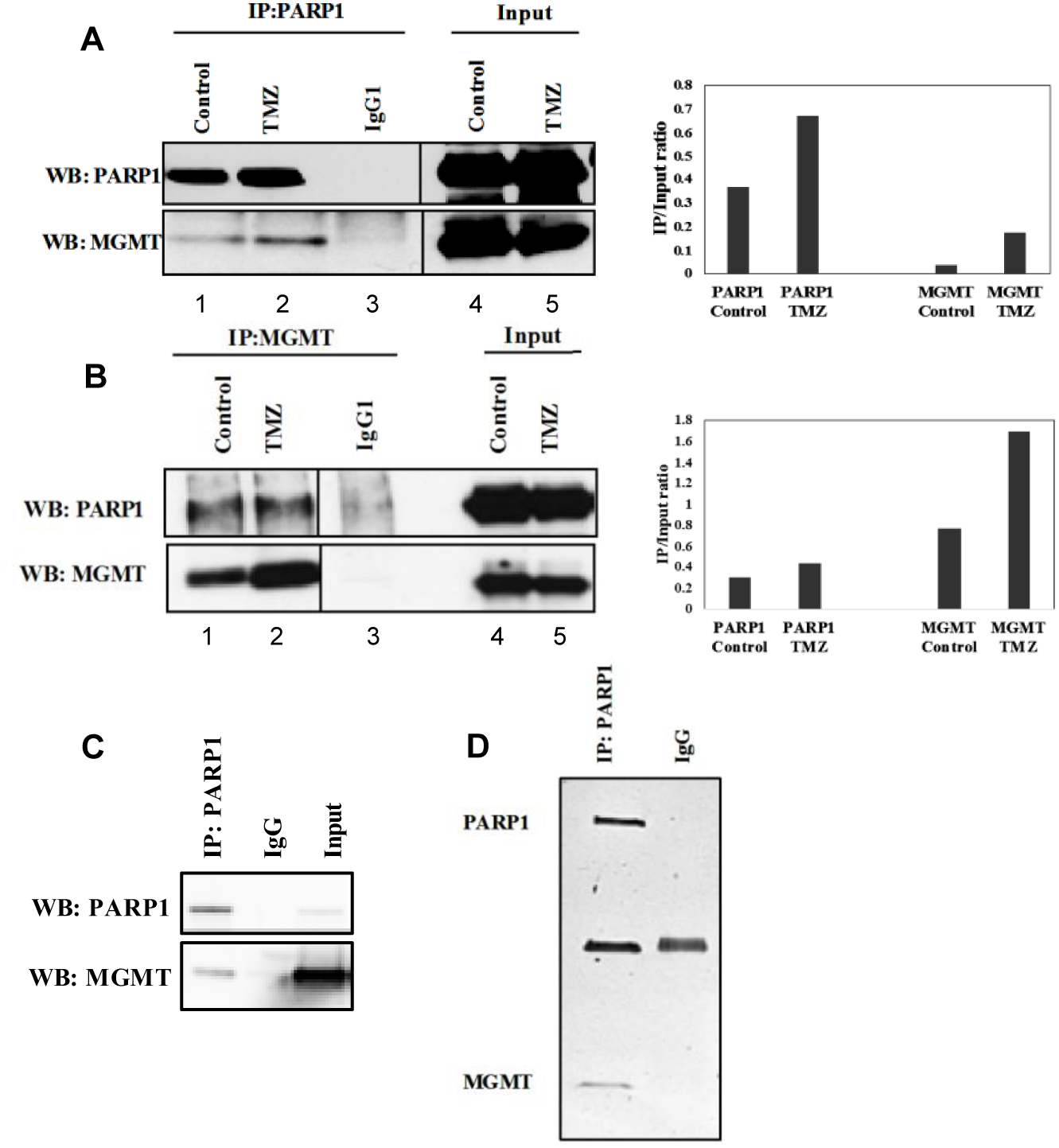
***A***, PARP1/MGMT direct interaction by co-IP. ***A***, EW-8 cells were treated with TMZ (1 mM, 2 hrs), PARP1 was pulled down with α-PARP1 and immunoblotted with α-PARP1 (top) or α-MGMT (bottom). IgG1 control is in the middle lane. Band quantitation is plotted as IP/Input ratio on the right. ***B***, EW-8 cells were treated with TMZ (1 mM, 2 hrs), MGMT was pulled down with α-MGMT and immunoblotted with α-PARP1 (top) or α-MGMT (bottom). IgG1 control is in the middle lane. Band quantitation is plotted as IP/Input ratio on the right. ***C***, Mixed purified PARP1 and MGMT proteins (1:1) were processed for co-IP. PARP1 was pulled down with α-PARP1 and immunoblotted with α-PARP1 (top) or α-MGMT (bottom). IgG control is in the middle lane. ***D***, SDS-PAGE and silver staining for samples prepared as in (C).

To test for direct interaction between PARP1 and MGMT, we carried out co-immunoprecipitation of purified full-length PARP1 and MGMT proteins (41) (**Fig. 3C**). An equal amount of PARP1 and MGMT proteins was mixed and PARP1 was precipitated with PARP1 and MGMT antibodies using protein G beads. Non-specific IgG (rabbit) or IgG1 (mouse) were used as negative control. SDS-PAGE analysis of the immunoprecipitated proteins followed by Western blots (**Fig.3C**) and silver staining (**Fig. 3D)** showed that PARP1 can interact with MGMT, confirming the direct physical interaction of these proteins.

We validated the direct protein-protein interaction using microscale thermophoresis and assessment of binding affinity (**Fig. 4**). The His-labeled purified PARP1 molecule was kept at a constant concentration (100 nM), while the concentration of the non-labeled binding partner MGMT varied between 13 nM and 21 μM. An average Kd = 165 nM was derived for the PARP1/MGMT interaction, confirming strong affinity of these proteins toward each other.

**Figure 4.**
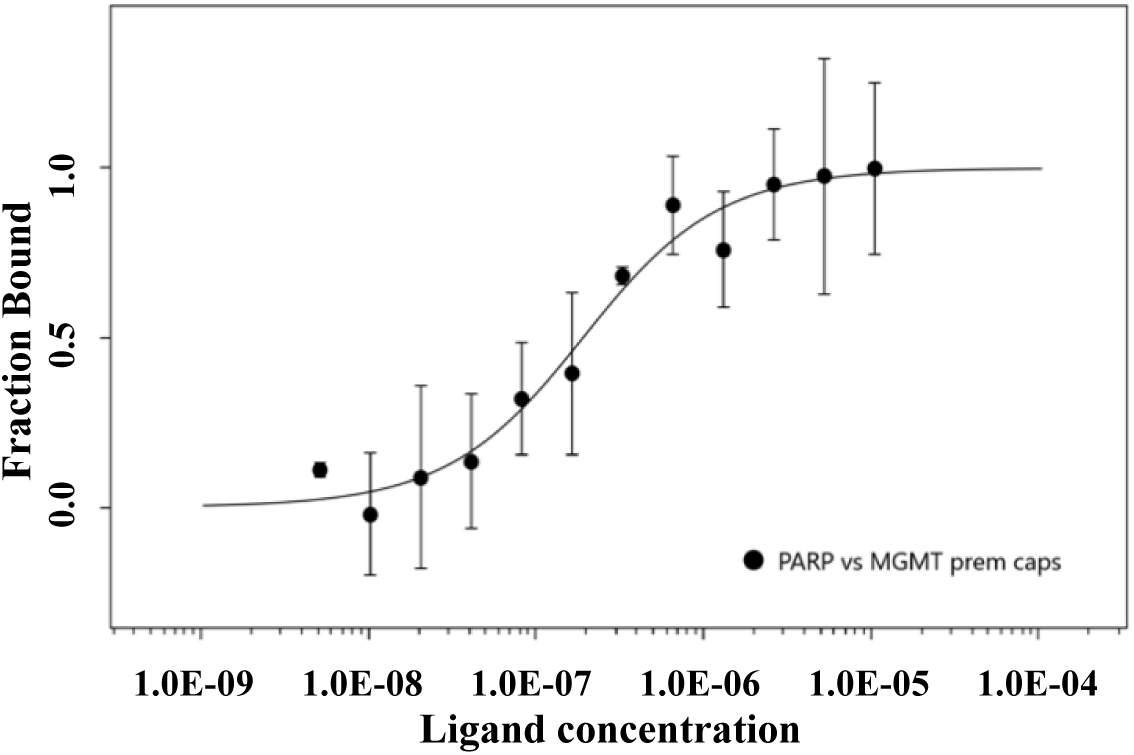
Microscale thermophoresis (MST) analysis of the PARP1/MGMT direct interaction. Recombinant PARP1 (His-labelled) and MGMT were used for the experiment. MST measurement was performed using the Monolith NT.115 at 17% LED power and medium MST power. An MST-on time of 10 sec was used for analysis, and an average Kd of 165 nM was derived for this interaction (n=3 independent measurements).

Confocal imaging analysis of immunostained proteins allows visualization of protein co-localization. For this purpose, the EW-8 cells ± TMZ treatment (1 mM for 2 hrs) were pre-incubated with the corresponding primary antibodies, the nuclei were stained with Hoeschst 33342, and PARP1 and MGMT were stained with Alexafluor 488 and Alexafluor 647, respectively (**Fig. 5**). The overlap of PARP1 (green) and MGMT (red) fluorophores in nuclei (blue) represented co-localization sites (white pixels) in the confocal images. The number of pixels in TMZ-treated cells increased (**Fig. 5B** and **D**), indicating that DNA damage induces co-localization of these proteins, consistent with our co-IP results in these cells (**Fig. 3A** and **B**).

**Figure 5.**
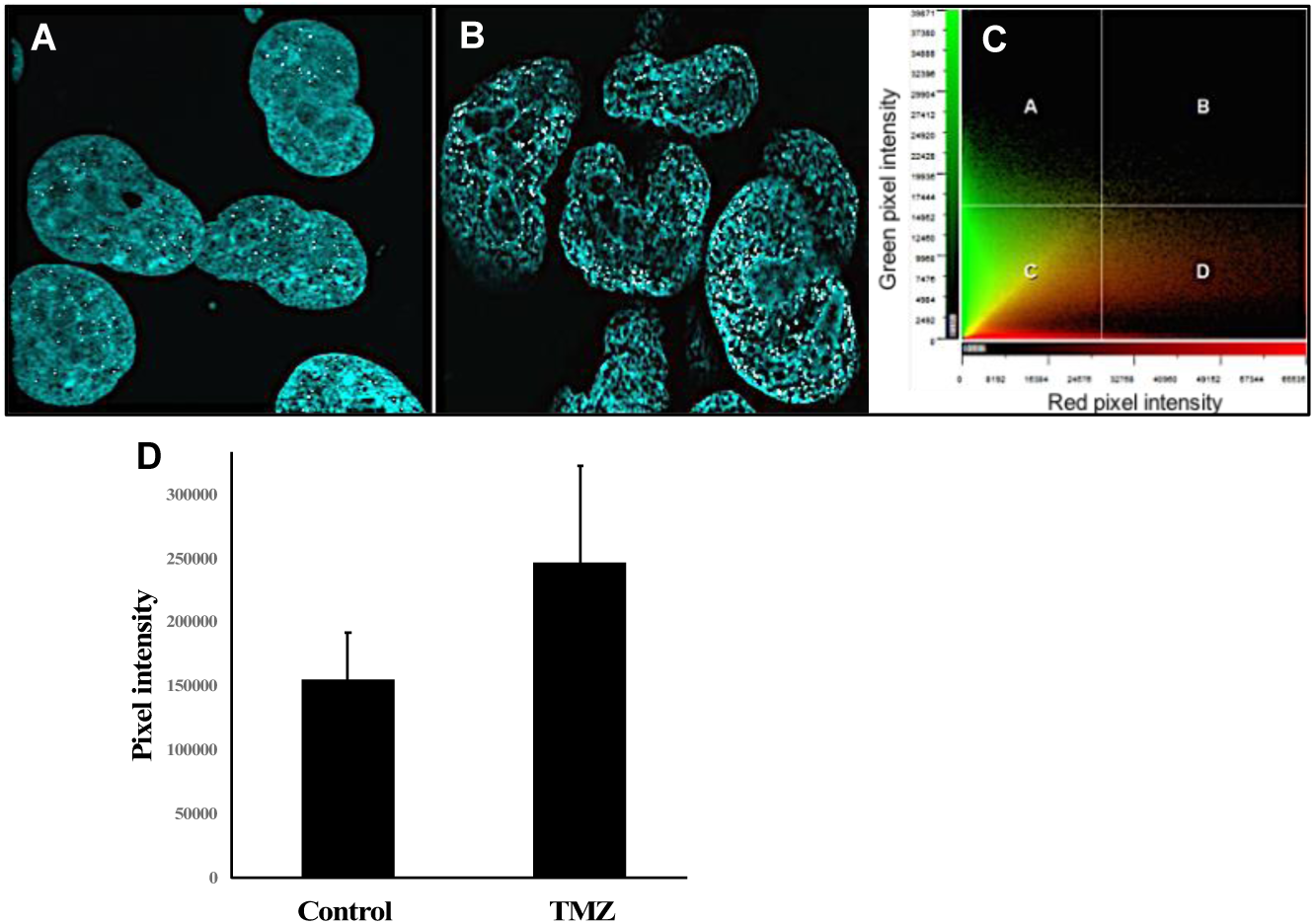
Confocal imaging analysis of PARP1 and MGMT proteins. EW-8 cell nuclei stained with Hoechst 33342 (blue). The white pixels indicate co-localization of PARP1 (green) and MGMT (red). ***A***, Control. ***B***, TMZ-treated cells (1mM, 2hrs). ***C***, Scatterplot of red and green pixel intensities, which result in white pixels upon overlap. Images were developed with Fluoview FV3000. Co-localization analysis was done using CellSens software (V2.1). ***D***, Quantification of co-localization in (A) vs (B) by identification of pixels with high intensity of staining for both fluorophores using co-localization algorithm built-in the CellSens software (V.2.1).

## Discussion

Two canonical DNA repair pathways are involved in repair of TMZ-induced DNA adducts: PARP1-dependent BER and MGMT pathways (MMR is involved when MGMT is inactive) (42). We have shown that inhibition of PARP1 and MGMT leads to the same degree of sensitization by TMZ, i.e. the removal of N3-meA, N7-meG, and O6-meG by PARP1 and MGMT is performed in an *epistatic* manner. This is the first description of PARP1/MGMT cooperation in DNA damage repair (33). Importantly, our cell survival data show that this cooperative relationship is independent of both the intrinsic sensitivity of Ewing sarcoma cells to TLZ and TMZ, and the DNA trapping potency of the PARP1 inhibitor tested.

The epistatic mechanism of DNA repair by PARP1 and MGMT we describe suggests direct interaction between these proteins, which we validated both in cell extracts and with purified proteins. Only limited evidence from other laboratories suggests that MGMT may interact with other proteins and have other functions (43,44). Cells not exposed to any DNA-damaging agent possess smaller amounts of the PARP1/MGMT complex, while induction of DNA damage leads to increased complex assembly (**Figs. 3A** and **B**). This result may indicate that DNA damage induces tight association of MGMT with PARP1 to enhance repair of the methyl adducts.

Consistent with our co-immunoprecipitation data, confocal imaging of immunostained PARP1 and MGMT proteins showed co-localization in EW-8 cells, and increased co-localized foci in response to DNA damage. Our co-immunoprecipitation with purified proteins and microscale thermophoresis analysis results indicate that neither DNA nor PARylation of MGMT by PARP1 are required for physical interaction.

Thus, PARP1/MGMT interaction may enhance repair when DNA damage is induced, suggesting that this interaction could be more efficient than PARP1 or MGMT repair processes canonically known to be independent. Further investigation of structural aspects of this novel interaction is underway to provide evidence for developing chemical agents to disrupt the PARP1/MGMT complex, which may be efficacious therapeutics for Ewing sarcoma and other cancers dependent on BER or repair by MGMT.

## Experimental procedures

### Cell culture and proliferation

Ewing sarcoma ES-4, ES-6, ES-7, and EW-8 cell lines were cultured in RPMI-1640 medium (*HyClone*) supplemented with 10% heat-inactivated FBS (*Millipore Sigma*). Cells were maintained at 37°C in a humidified atmosphere with 5% CO_2_.

The Alamar Blue® assay was used to assess cell viability (*BioRad*). Cells were seeded to reach 20-40% confluency (lower confluency for rapidly growing cells). TLZ, TMZ, and O6-benzylguanine were added to wells 24 hrs after cell seeding, and incubated for 96 hrs. After 2 hrs incubation of cells in 24-well plates (1,000 μl of culture medium), 10% v/v Alamar Blue (100 μl) was added and fluorescence was measured (excitation 530 nm, emission 590 nm). Wells containing RPMI 1640 (*Hyclone*), 10% FBS (*Millipore Sigma*) and untreated cells, and 10% v/v Alamar blue were used as positive controls. Wells with culture medium without cells containing 10% v/v Alamar Blue were negative controls. Fluorescence was recorded on a Spectra Max plate reader, using the Alamar Blue protocol provided by *Softmax Software*. Statistical analyses and curve plotting (4-parameter polynomial analysis) were performed using standard equations included in the GraphPad Prism 7.0c package (*GraphPad Software Inc., USA*).

### Protein expression and purification

Plasmids for PARP1 protein constructs were provided by John Pascal (41). Expression and purification were performed following the protocol described elsewhere with minor modifications (41). Full-length His6-PARP1was overexpressed using *Escherichia coli* strain BLR (DE3) pRARE *(Novagen)*. Transformed cells were grown in 2 x LB medium and protein was induced with 0.2 mM isopropyl ß-D-thiogalactopyranoside and 0.1 mM of ZnCl_2_ at 16°C for 20 hrs. Cell lysate was prepared by sonication and clarified by ultracentrifugation. His6-PARP1 protein was purified with 2 ml Ni-NTA agarose *(Qiagen)*, 1 ml HiTrap Heparin HP *(GE Healthcare)*, and HiTrap SP HP *(GE Healthcare)*. Full-length His6-MGMT (45) was overexpressed using *Escherichia coli* strain BLR (DE3) pRARE *(Novagen)*. Transformed cells were grown in 2xLB medium and His6-MGMT was induced with 0.5 mM IPTG at 16°C for 20 hrs. Cell lysate was prepared by sonication and clarified by ultracentrifugation. His6-MGMT protein was purified with 3 ml Ni-NTA agarose *(Qiagen)* and 1 ml Source S *(GE Healthcare)* steps.

### Co-immunoprecipitation

Cells were grown to near confluency in 10 cm dishes. Whole cell lysates were prepared using IP lysis buffer (*Pierce*) supplemented with Halt protease and phosphatase inhibitor (*Thermo Fisher Scientific*) plus PMSF (*Millipore Sigma*), according to standard protocols. Co-immunoprecipitation was done with endogenous and purified PARP1 and MGMT proteins. For purified proteins, the constructs were mixed 1:1 (2 µg total protein concentration each). Pull-downs were performed using SureBeads protein G magnetic beads (*BioRad*) according to the manufacturer’s protocol. Bound complexes were eluted with Invitrogen NuPage loading buffer (*Invitrogen*) and then evaluated by Western blotting. Antibodies used for pull-down include MGMT (*Santa Cruz Biotechnology*), PARP1 (*Cell Signaling Technology)*, and IgG/IgG1 (*Cell Signaling Technology*).

### Immunoblotting

Cells were lysed using RIPA buffer (*89900, Pierce*) according to standard protocols. Samples were separated on a 4-12% gradient gel (*NP0321, Invitrogen*) and then transferred onto a PVDF or nitrocellulose membrane. Membranes were blocked with 3% BSA in TBS-T for 1 hr at room temperature, then incubated with primary antibody overnight. After secondary antibody incubation and washing, membranes were developed using enhanced chemiluminescence (*PerkinElmer*). Antibodies used include PARP1 (*Cell Signaling Technology*) and MGMT (*Santa Cruz Biotechnology*).

### Immunostaining and confocal imaging

1×10_5_ of EW-8 cells were plated into 24-well plates with glass coverslips in sterile conditions and incubated for 48 hrs at 37°C for cells to settle. Cells were treated with 1 mM of TMZ for 2 hrs, and then fixed with 4% formaldehyde for 15 min at room temperature. After blocking for 1 hr with blocking buffer (1xPBS/5% normal goat serum/0.3% Triton x-100) cells were incubated with primary antibodies (PARP1, 1:1600, *Cell Signaling Technology*; MGMT, 1:50, *Santa Cruz Biotechnology*) overnight at 4°C on a cold room shaker. The following day, cell nuclei were stained with Hoechst 33342 (blue) and incubated with fluorochrome-conjugated secondary antibodies (PARP1, Alexafluor 488 – green and MGMT, Alexafluor 647 – red, *Abcam*) for 2 hrs at room temperature. Images were captured with an Olympus Ix80 microscope at 100 X magnification. Images (Z-stacks) were developed and analyzed with Fluoview and CellSens software (v2.1) to identify pixels with a high degree of co-localization (high intensity of staining for both fluorophores) using the built-in co-localization algorithm.

### Microscale Thermophoresis

PARP1 protein (*Trevigen*) was labeled with His using the Protein Labeling Kit RED-NHS 2_nd_ Generation (*NanoTemper Technologies*). The labeling reaction was performed according to the manufacturer’s instructions using the labeling buffer provided in the kit, applying a concentration of 2.7 μM protein (molar dye: protein ratio ≈ 5:1) at room temperature for 30 min in the dark. Unreacted dye was removed with the dye removal column included in the kit, and equilibrated with PBS (pH 7.5). The labeled protein PARP1 was adjusted to 100 nM with PBS supplemented with 0.005% of Tween-20. The ligand MGMT (*Novus Biologicals*) was dissolved in PBS supplemented with 0.005% of Tween-20, and a series of twelve 1:1 dilutions were prepared using the same buffer, producing ligand concentrations ranging from 13 nM to 21 μM. For measurement, each ligand dilution was mixed with one volume of labeled PARP1, yielding a final concentration of PARP1 of 50 nM and final ligand concentrations of 6.5 nM to 10.5 μM. After 20 min, the samples were loaded into Monolith NT.Automated Premium Capillary Chips (*NanoTemper Technologies*). Microscale thermophoresis was measured using a Monolith NT.Automated instrument (*NanoTemper Technologies*) at an ambient temperature of 25°C. Instrument parameters were adjusted to 17% excitation and medium power. Data from 3 independently pipetted measurements were analyzed (MO.Affinity Analysis software version 2.3, *NanoTemper Technologies*) using the signal from an instrument-on time of 10 s.

## Acknowledgements

We are grateful to our colleague Patrick Sung for advice and for editing our manuscript (Youngho Kwon is supported by NCI grant R35 CA241801 awarded to Patrick Sung). We would also like to thank Dinorah Leyva at *NanoTemper* for her expertise and help with the MST analysis, and our colleague Peter Houghton for thoughtful discussions.

## Conflict of interest

The authors declare that they have no conflicts of interest with the contents of this article.

## Author contributions

RK, PS, and YK designed the research project; DA, YK, JC, and RL performed experiments; RK, PS, and YK analyzed data; RK wrote the manuscript; and YK, PS, and DA edited the manuscript.

## FOOTNOTES

Funding was provided by the Greehey Children’s Cancer Research Institute at The University of Texas Health Science Center at San Antonio (JC, RL, DA, RK); an Emerging Scientist Award from the Childhood Cancer Research Fund (DA, RK); and NCI R35 CA241801 (YK).

## The abbreviations used are

TLZ: (talazoparib);
TMZ: (temozolomide);
PARP1: (Poly(ADPribose) Polymerase I);
MGMT: (O6-methylguanine methyltransferase);
PPTP: (Pediatric Preclinical Testing Program).

